# Opposite Priming Effects on Identity vs. Category Recognition Require Conscious Awareness

**DOI:** 10.1101/2025.09.13.676001

**Authors:** Samaneh Navab Kashani, Reza Khosrowabadi, Mohammad-Reza A. Dehaqani

**Affiliations:** Institute for Cognitive and Brain Sciences, Tehran, Iran; College of Engineering, School of Electrical and Computer Engineering, University of Tehran, Tehran, Iran; School of Cognitive Sciences, Institute for Research in Fundamental Sciences (IPM), Tehran, Iran

**Keywords:** Conscious priming, Unconscious priming, Visual perception, Identity recognition, Category recognition, Binocular rivalry, Drift Diffusion Model

## Abstract

How conscious and unconscious priming differentially modulate object recognition at different levels of abstraction (identity vs. category) remains incompletely understood, despite extensive research. We used a binocular rivalry paradigm with Continuous Flash Suppression (CFS) to manipulate conscious awareness with image or word primes while participants performed a name-picture verification task probing identity and category recognition for faces and animal bodies. Behavioral results revealed a striking dissociation: consciously perceived primes facilitated identity recognition but impaired category recognition. This effect was most pronounced with image primes, face targets, and right visual field presentations. Under unconscious priming, no such effects were observed. To elucidate the underlying mechanism, we applied a Drift Diffusion Model, which revealed that conscious priming selectively enhanced the efficiency of evidence accumulation and introduced a pre-decisional bias for identity decisions, with no reliable change for category decisions. Our findings demonstrate a double dissociation where conscious awareness is required for priming to exert robust and opposite effects on identity and category recognition. This finding challenges the view of priming as a uniformly facilitatory process, providing a new mechanistic framework for understanding how consciousness and abstraction level interact to shape visual perception.

**Highlights:**

- Conscious awareness is required for priming to exert robust effects on recognition
- The priming aftereffects can be facilitatory or impairing depending on the level of recognition
- Conscious priming facilitated identity recognition but impaired category recognition.
- Dissociation of identity and category by conscious priming were strongest for image (vs. word) primes, face (vs. animal) targets, and right (vs. left) visual field presentations.
- Drift diffusion modeling revealed increased drift rate and pre-decisional bias, for identity but not for category decisions in conscious priming.

## Introduction

Experiments on priming have played a pivotal role in advancing theories of visual perception, demonstrating reliable effects at both the behavioral level (Jiang et al., 2007; Marsolek, 1999; Stein et al., 2020) and the neural level (Balconi, 2006; Dehaene et al., 1998; Diaz & McCarthy, 2007; Fang & He, 2005), and establishing priming as a powerful tool for probing the mechanisms by which the brain encodes and interprets visual information (e.g. facilitated responses and modulated neural activation for repeated or related items). Yet, how conscious and unconscious priming differentially modify identity and category recognition remains poorly understood (Amihai et al., 2011; Chien et al., 2023). Experimental (Breitmeyer et al., 2005; Dehaene et al., 2001; Kouider & Dehaene, 2007; Pessoa et al., 2005; Weibel et al., 2013) and neurophysiological (Berti & Rizzolatti, 1992; Trevethan et al., 2007) evidence indicates that unconscious primes can modulate subsequent perceptual and semantic processing (Kiefer & Spitzer, 2000; Kouider & Dehaene, 2007), however identity recognition often demands consciousness (Amihai et al., 2011; Moradi et al., 2005).

Research on priming has long established that even stimuli outside of awareness can influence perception and cognition, but the depth of this influence depends on conscious access. Recent work by Chien et al. found that unconscious semantic priming occurs only for directly associated concepts, whereas higher-level or abstract associations require conscious awareness (Chien et al., 2023). In the domain of face recognition, there is no strong evidence that an individual’s identity or attributes can be extracted without awareness (Amihai et al., 2011). Conscious and unconscious priming differ markedly in scope: unconscious primes evoke fragmentary, limited influences, while conscious primes can robustly modulate detailed recognition and semantic integration (Amihai et al., 2011; Chien et al., 2023; Ortells et al., 2006). However, despite extensive priming research, how awareness state alters the specific impact of priming on object recognition at different levels (identity vs. category) remains only partially understood.

Identity and category recognition engage distinct cognitive and neural processes, however their modulation by priming remains underexplored. Recognition organizes objects into superordinate (e.g., animal), basic (e.g., dog), and subordinate (e.g., Bulldog) levels, with basic categories processed most rapidly (Johnson & Mervis, 1997; Rosch et al., 1976). Neuroscience reveals faster category encoding in the inferior temporal cortex compared to identity (Dehaqani et al., 2016), suggesting distinct pathways—feedforward for categorization and feedback for identification. This dissociation suggests that initial feedforward activity is sufficient for rapid categorical basic recognition, whereas pinpointing an exact identity engages recurrent, top-down mechanisms commonly linked to conscious awareness (Dehaqani et al., 2016; Koivisto & Rientamo, 2016).While this framework differentiates recognition levels, how priming alters these dynamics across consciousness states remains unclear, necessitating a targeted investigation.

To address this, we employed a binocular rivalry setup—presenting word or image primes to one visual field, suppressed by a rival stimulus—offering precise control over consciousness, unlike traditional masking methods. Paired with a name-picture verification task using faces and animal bodies as targets, this approach tests priming across target stimuli and prime types. Prior studies often treat these processes in isolation, overlooking their interplay across conscious states and prime types. For example, research on word versus image primes has shown that image primes are generally more effective for identification, while pictures provide faster access to conceptual meaning than words (Carr et al., 1982; Potter & Faulconer, 1975; Reinitz et al., 1989). Similarly, studies focusing on faces have highlighted the critical role of consciousness in identity recognition, with unconscious priming producing at best limited effects on coarse categorical judgments such as gender or race (Amihai et al., 2011; Axelrod & Rees, 2014; Moradi et al., 2005). While these findings provide valuable insights within specific domains, they leave unresolved how priming effects generalize across different prime modalities and target categories under varying states of awareness, a gap the present study aims to address.

This study tackles this gap, probing the scope and mechanisms of priming effects under controlled conditions, with a novel focus on these variables. Drift Diffusion Model (DDM) analysis further enhances our method, quantifying contributions of consciousness, level of recognition, and relevant prime compared to irrelevant prime in a unified framework, while also enabling us to uncover the underlying cognitive mechanisms and latent aspects of behavior that remain invisible in traditional accuracy and reaction-time measures.

## Material and methods

### Participants

Twenty-two healthy, right-handed volunteers (mean age = 35.7 years, range = 32–43 years, 8 females) with no prior experience with the task participated in the experiment. All had normal or corrected-to-normal vision and provided written informed consent.

### Stimuli and apparatus

Participants were seated in a darkened room, facing a monitor (1920 × 1080 resolution, 144 Hz refresh rate, 1 ms response time) to minimize stimulus aftereffects. The viewing distance was fixed at 64 cm, and stimuli were viewed through an adjustable mirror stereoscope mounted on a chin rest, with presentations set against a gray background.

Stimuli consisted of Persian words and grayscale images representing four classes: women, men, cats, and dogs, tested at two recognition levels (identification and categorization). As shown in Figure S1 in Supplementary material, each class included a category name and symbolic image denoting the category (woman, man, cat, dog), and three identities presented as both target name/picture and image/word prime (e.g., women: Laleh, Minou, Afarin; men: Farshad, Pedram, Bijhan; cats: Persian, Bengal, Brown; dogs: Bulldog, Shepherd, Doberman). To induce suppression (Tsuchiya et al., 2009), Continuous Flash Suppression (CFS) patterns were generated using the MATLAB Psychtoolbox (Brainard, 1997), featuring Mondrian noise—colored squares of varying sizes flashed at 20 Hz.

Face and animal images, sourced from Google searches, along with word stimuli and category symbols, were luminance-matched using the SHINE Toolbox (see Figure S2 in Supplementary Material). All stimuli were presented via the MATLAB Psychtoolbox. Except for prime stimuli (4° visual angle), all stimuli and CFS patterns subtended 6° of visual angle to avoid retinotopic overlap (MI. Posner & Y Cohen, 1980; Posner & Cohen, 1984). Non-prime stimuli were framed with a textured black-and-white border extending to 6.5° visual angle (see Figure 1A).

**Figure 1:**
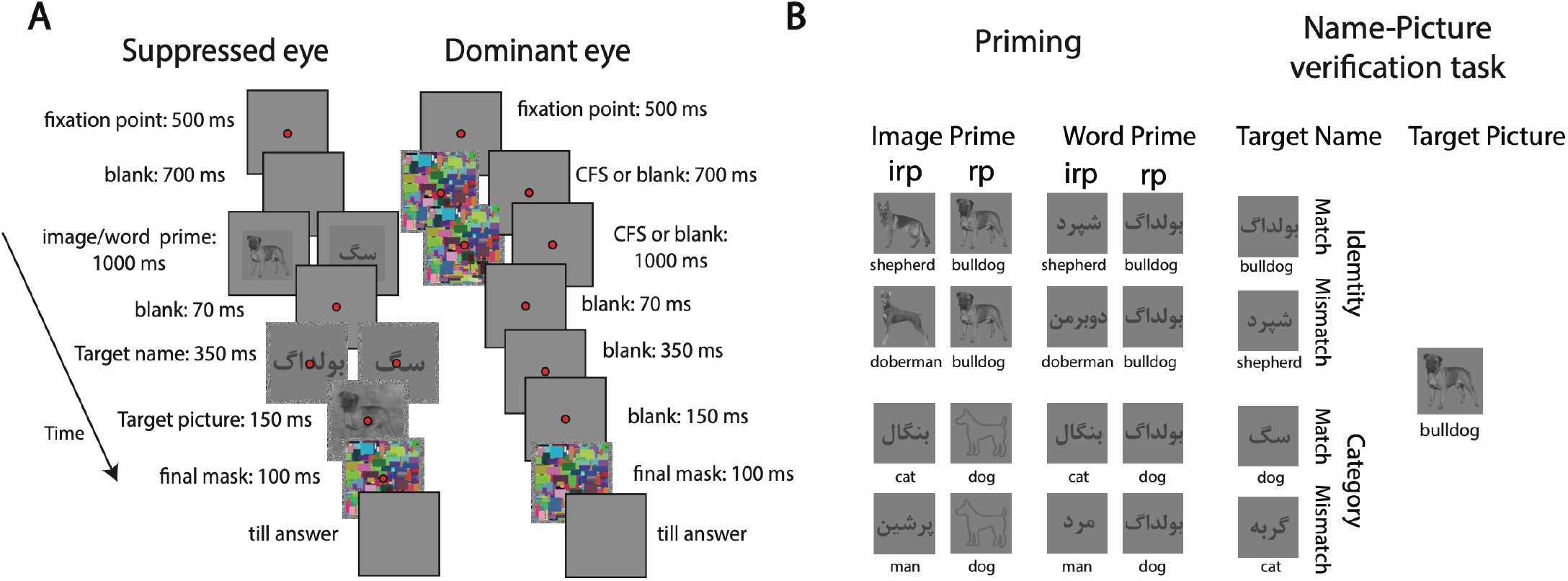
Experimental paradigm. **(A)** Schematic of the binocular rivalry task with Continuous Flash Suppression (CFS). The left frame is presented to the left /right visual field (suppressed eye), and the right frame to the right/left visual field (dominant eye), depicting conscious (no CFS) and unconscious (CFS) conditions. Prime type (words or images) is shown at identity or category levels, followed by the name-picture verification task. **(B)** Stimuli in prime and the name-picture verification task. Each target picture is paired with four different target names: identity (e.g., “Bulldog”/“Shepherd”) or category (e.g., “Dog”/“Cat”), matching or mismatching the picture. Trials include four different prime: relevant image prime (rp; image), irrelevant image prime (irp; image), relevant word prime (rp; world), or irrelevant word prime (irp; word), varying by recognition level and matchness.

### Procedure and design

#### Binocular rivalry

To manipulate consciousness, we utilized binocular rivalry combined with Continuous Flash Suppression (CFS), a method superior to traditional masking (e.g., backward or forward masking) due to its prolonged subliminal presentation and precise control over perception (Axelrod et al., 2015; Tsuchiya & Koch, 2005). In binocular rivalry, as shown in Figure 1A, distinct stimuli are presented simultaneously to each visual field, with one consciously perceived while the other remains subliminal yet influential (Blake, 2001; Fang & He, 2005). Here, CFS employed Mondrian noise (colored squares flashing at 20 Hz) as a suppressor, dominating perception for extended periods (up to 50 s), while prime stimuli in the suppressed eye exerted unconscious effects (Tsuchiya & Koch, 2005). This setup, unlike breaking CFS (b-CFS) paradigms (Jiang et al., 2007), allowed us to assess subliminal priming’s impact on a subsequent task rather than suppression breakthrough. Additionally, presenting primes foveally but lateralized to one visual field enabled the exploration of hemispheric asymmetries, one of the factors manipulated in our design.

#### Name-picture verification task

We adapted a name-picture verification task (Collin & Mcmullen, 2005) to probe identity and category recognition under priming. Participants judged whether a name target matched a subsequent picture, with match and mismatch trials balanced (50% each). Illustrated in figure 1B, identity trials tested subordinate-level recognition (e.g., “Bulldog or Shepherd” target name followed by a Bulldog picture), while category trials assessed basic-level recognition (e.g., “Dog or Cat” target name followed by a Dog picture). Name-picture verification task controlled the processing level by presenting the name target first, ensuring decisions reflected the specified recognition level. Mismatch trials used within-category distractors for identity (e.g., Bulldog-Doberman) and cross-category distractors for basic level or categorization task (e.g., Bulldog-Cat), avoiding superordinate overlap. Primes—either words or images—preceded the task, with relevant primes matching the target name and irrelevant primes drawn from unrelated categories (e.g., “Doberman” for a Bulldog-Shepherd trial) or superordinate levels (e.g., “Man” for a Cat-Bulldog trial), as shown in Figure 1B.

#### Experimental design

Participants first completed a training phase without stereoscopes or primes, identifying men, women, cats, and dogs via paper materials and practice blocks of the name-picture verification task (identity level only). A fixation point (0.25° visual angle) turned green for correct responses and red for errors, ensuring familiarity; only those achieving >95% accuracy proceeded. The main experiment used a within-participant design with no feedback, maintaining a red fixation point. Participants focused on this point and responded via right-hand keypresses (right key for match, left for mismatch) while viewing stimuli through the stereoscope, adjusted until a single central dot was visible.

In the main task, primes and name-picture target stimuli appeared in the left visual field for half the trials and the right visual field for the other half, testing hemispheric effects. In unconscious conditions, CFS noise suppressed primes in the opposite field (e.g., right visual field CFS for left visual field primes). Each trial began with a 500-ms red fixation point, followed by 1700-ms CFS (unconscious) or a blank screen (conscious). During the final 1000 ms, a prime (word or image) faded in (contrast: 0.15% to 0.4%). After a 70-ms blank, a name target appeared for 350 ms, followed by a 150-ms picture in a noisy phase (0.4% frequency), masked by 100-ms bilateral Mondrian noise. A gray screen persisted until response. Conscious trials omitted CFS, rendering primes visible for 1000 ms. Name targets and primes varied randomly across identity and category levels, with symbolic images (e.g., cat, dog) used as category level primes. The paradigm is detailed in Figure 1A.

To ensure that the primes presented under CFS remained entirely unconscious, participants were interviewed after each of the three scheduled breaks regarding their subjective visual experience. If there was any suspicion that the suppressed primes might have become visible, we either readjusted the stereoscope setup and re-administered the visibility test or, when this was not possible, excluded the participant from the study. In practice, one participant was excluded on this basis.

As shown in experimental paradigm (Figure 1), in this framework, we have 7 pairs of conditions, and each trial is a combination of these 7 conditions, resulting in a response time (RT) and performance of the subject: Priming (relevant prime/irrelevant prime), Consciousness (conscious prime/unconscious prime), Recognition level (identity/category), Prime type (image/word), Target stimuli (face/animal), Laterality (left/right visual field), Matchness (match/mismatch). With six stimulus samples in the target set (three women, three men, three cats, and three dogs), participants responded to 768 trials in total.

### Statistical analysis

Reaction time (RT) and accuracy data were recorded for each trial. Only correct responses with RTs between 50 ms and 6000 ms were analyzed, excluding <5% of trials. Mean RTs were computed per participant, with RT as the primary focus due to its sensitivity to priming effects.

Two-tailed Wilcoxon signed-rank tests assessed effect reliability via planned pairwise comparisons, while Pearson’s correlation coefficients evaluated relationships between variables. P-values and effect sizes are reported to three decimal places, with ns ≥ 0.05; * < 0.05; ** < 0.01; ***<0.001. The p-values were corrected using the Bonferroni method for multiple hypothesis testing.

### Prime Index (PI)

To quantify priming effects across conditions, we defined a Prime Index (PI) as the difference in mean RT between trials with a relevant prime (task-related) and a baseline condition with an irrelevant prime (unrelated identity or category level). Both relevant and irrelevant prime shared the same recognition level (identity or category), ensuring level-specific comparisons. PI served as a standardized metric to compare conscious vs. unconscious, left vs. right visual field, face vs. animal, and word vs. image prime conditions.

### Drift Diffusion Modeling (DDM)

To capture how participants accumulated evidence over time to make perceptual decisions, we applied the Drift Diffusion Model (DDM) (Ratcliff, 1978; Ratcliff et al., 2016; Ratcliff & McKoon, 2008). The DDM assumes that decisions are based on a sequential accumulation of noisy sensory evidence, which continues until the accumulated evidence reaches one of two boundaries, triggering a response. We model (DDM) to characterize how evidence was accumulated and transformed into perceptual decisions in the name–picture verification task. Within this framework, the drift rate (*υ*) reflects the efficiency of evidence accumulation, such that higher drift corresponds to faster and more accurate discrimination between correct and incorrect response to name–picture pairs; in our task, drift is expected to vary with informative evidence of the prime and consciousness factors. The boundary separation (a) captures the level of response caution, with larger values indicating that participants required more evidence before making a decision, thus trading speed for accuracy. The starting point (z) represents a bias toward one response alternative that is influenced by prior expectations and tendencies, which in the context of our task may reflect a predisposition toward “correct” versus “incorrect” answer induced by priming. Finally, the non-decision time (*T*_*er*_) accounts for processes outside of evidence accumulation, including the perceptual encoding of stimuli under binocular rivalry and the motor execution of the response. By estimating these parameters separately across priming, consciousness condition, and level of recognition, we were able to test how conscious and unconscious primes influenced the dynamics of decision formation. Conceptually, the decision variable evolves as a noisy accumulation process:

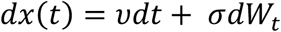

Where *υ* is the drift rate, representing the average rate of evidence accumulation toward the correct boundary, *σ* is the diffusion constant (set to 1 without loss of generality), and *dW*_*t*_ is a Wiener process capturing moment-to-moment noise in accumulation.

Decisions are made when the process first crosses one of two absorbing boundaries at 0 or a. The predicted response time is then the sum of the decision time and the non-decision component:

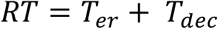

### Parameter fitting

For each subject and condition (priming × consciousness × recognition level), we fit the four parameters of DDM by maximizing the likelihood of observed response and RTs. The probability of each response was approximated using a logistic function, and the response time distribution was captured using a Gaussian approximation centered on the expected reaction time. This simplified likelihood approach has been used in prior applications of DDM when full analytic solutions are computationally expensive (Voss et al., 2013).

Optimization was implemented in MATLAB using constrained nonlinear minimization (fmincon), with appropriate parameter bounds to ensure meaningful estimates (e.g., non-decision times > 0, drift rates within a reasonable range). Parameter estimation was performed separately for each subject and condition to allow comparison across experimental manipulations.

## Results

We investigated the differential effects of conscious and unconscious priming on identity and category recognition using a binocular rivalry paradigm (Figure 1A). Participants completed a name-picture verification task, where word or image primes, preceded target faces or animal bodies, presented to the left or right visual field. Using these different conditions, namely priming (relevant/irrelevant prime), consciousness (conscious/unconscious prime), recognition level (identity/category), prime type (image/word), target stimulus (face/animal), and laterality (left/right visual field), we want to address all dimensions of the priming aftereffect. This design enabled us to assess how priming modulates recognition processes across conscious prime and unconscious prime.

### Robustness of basic-level advantage in conscious and unconscious priming

Extensive research highlights a basic-level advantage in object recognition, with faster and more accurate responses compared to subordinate levels (Fan et al., 2012; Jolicoeur et al., 1984; Rosch et al., 1976; Tanaka & Taylor, 1991). While well-documented in naming and categorization tasks, this effect is less explored in priming paradigms. Here, we observed a robust basic-level (category) advantage in both conscious and unconscious conditions, evident in performance and reaction time (RT) across relevant and irrelevant prime trials.

Performance was superior for category tasks compared to identity tasks under conscious (Figure 2A: *Performance*_*cat*−*id*_ = −0.038 ± 0.009, p = 0.002) and unconscious (Figure 2A: *Performance*_*cat*−*id*_ = −0.051 ± 0.012, p = 0.001) priming. This advantage held for 81% of participants (18/22; Figure 2B). Similarly, RTs were shorter for category tasks than identity tasks in conscious (Figure 2C: *RT*_*cat*−*id*_ = −353.5 ± 33.9 ms, p < 0.001) and unconscious (Figure 2C: *RT*_*cat*−*id*_ = −367.3 ± 35.0 ms, p < 0.001) conditions, consistent across all participants (Figure 2D). Each scatter plot point represents one participant, with error bars denoting standard errors of the mean (SEMs). These findings extend the basic-level advantage to priming contexts, aligning with prior work (Rosch et al., 1976).

**Figure 2:**
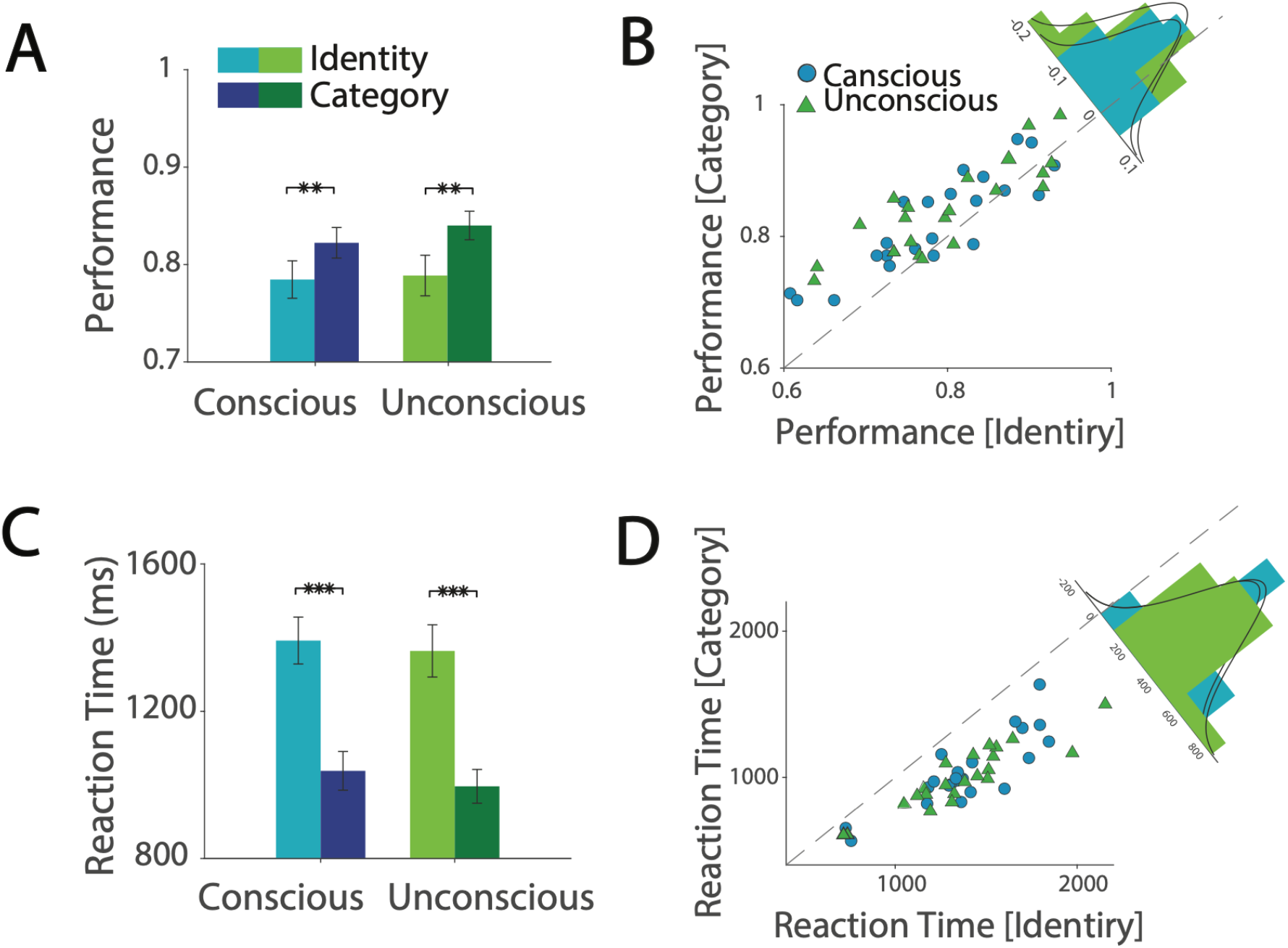
Basic level (category) advantage in priming. Panels **(A, B)** show accuracy, and **(C, D)** show reaction times for conscious and unconscious priming at identity and category levels. A basic-level (category) advantage is evident in faster and more accurate responses, modulated by consciousness. Histograms on the diagonal depict differences in performance and reaction time between category and identity trials. Error bars represent standard errors of the means (SEMs) across participants. Asterisks denote two-tailed tests: *\*\* < 0.01; *** < 0.001*.

Replicating the basic-level advantage provided a stable baseline from which to assess the novel priming effects. This replication not only confirmed a robust and well-established phenomenon in object recognition but also ensured that any deviations induced by priming could be interpreted against a reliable benchmark. Having established this foundation, we next examined whether conscious and unconscious primes differentially modulate response at the identity versus category level, revealing a dissociation that has not been documented in prior research.

### Dissociable effects of conscious priming on identity and category recognition

Our primary aim was to determine whether priming differentially affects identity and category recognition under conscious and unconscious conditions in a binocular rivalry paradigm. We measured participants’ reaction times (RTs) in a name-picture verification task following word or image primes and computed prime index (PI) as the mean RT difference between relevant and irrelevant prime trials. Under conscious priming, a striking dissociation emerged: identity recognition was facilitated (Figure 3A: *PI*_*id*_ = −104.5 ± 38.4 ms, p = 0.015), with faster RTs to relevant primes compared to irrelevant prime, indicating positive priming. In contrast, category recognition was impaired (Figure 3A: *PI*_*cat*_ = 57.7 ± 20.4 ms, p = 0.028), with slower RTs relative to irrelevant primes. This divergence was significant (Figure 3A: Δ*PI*_*cat*−*id*_ = 162.2 ± 42.7 ms, p = 0.002), and appear in near all participants (Figure 3B: 18/22). highlighting opposing priming effects within conscious conditions. No such dissociation appeared under unconscious priming, suggesting these effects are conscious-dependent.

**Figure 3:**
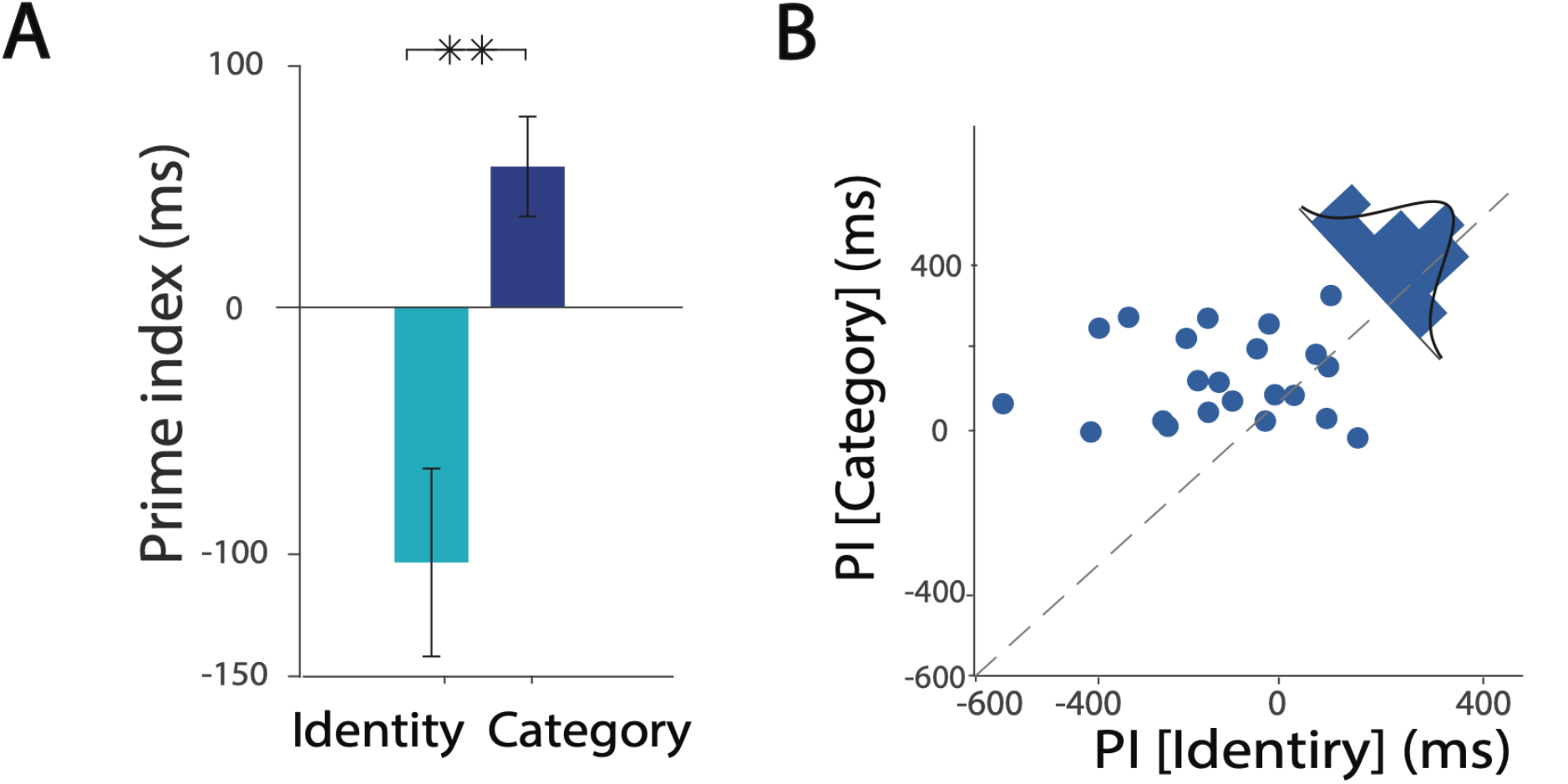
Dissociable conscious priming effects on identity and category recognition. **(A)** Prime Index (PI), calculated as the mean reaction time (RT) difference between relevant and irrelevant prime trials, varies by recognition level under conscious priming. Identity recognition shows facilitation (faster RTs to relevant compared to irrelevant primes; *PI*_*id*_ = −104.5 ± 38.4 ms, p = 0.015), while category recognition exhibits impairment (slower RTs to relevant compare to irrelevant primes; *PI*_*cat*_ = 57.7 ± 20.4 ms, p = 0.028). **(B)** The scatter plot and histogram depict inter-subject variability, with most participants (18/22) showing faster identity RTs and slower category RTs under conscious conditions. Asterisks denote two-tailed tests: *\*\* < 0.01*.

To elucidate the condition dependency underlying priming, and offering insights into the underlying brain systems, we manipulated six factors: priming (relevant vs. irrelevant prime), recognition level (identity vs. category), prime type (word vs. image), consciousness (conscious vs. unconscious), laterality (left vs. right visual field), and target stimuli (face vs. animal). This multivariate design, detailed in Figure 1B, allowed us to examine how these variables modulate visual recognition and using the PI metric to quantify condition-specific effects and their interactions.

### Influence of prime type, laterality, and target stimuli on priming effects

To further dissect the dissociation between identity and category recognition, we analyzed the roles of prime type (image vs. word), laterality (left vs. right visual field), and target stimuli (face vs. animal) under conscious priming. Prior studies suggest that image and word primes influence recognition differently (Kiefer & Spitzer, 2000; Leopold et al., 2001), but their specific effects in a name-picture verification task remain unclear. As shown in Figure 4, image primes elicited a robust dissociation (Figure 4; image: Δ*PI*_*cat*−*id*_ = 293.5 ± 61.8 ms, *p*_*Bonf*_ < 0.01), with facilitation for identity and impairment for category recognition. In contrast, word primes showed no significant effect (Figure 4; word: Δ*PI*_*cat*−*id*_ = 25.3 ± 64.3 ms, *p*_*Bonf*_ ≈ 1), failing to differentiate the two levels.

**Figure 4:**
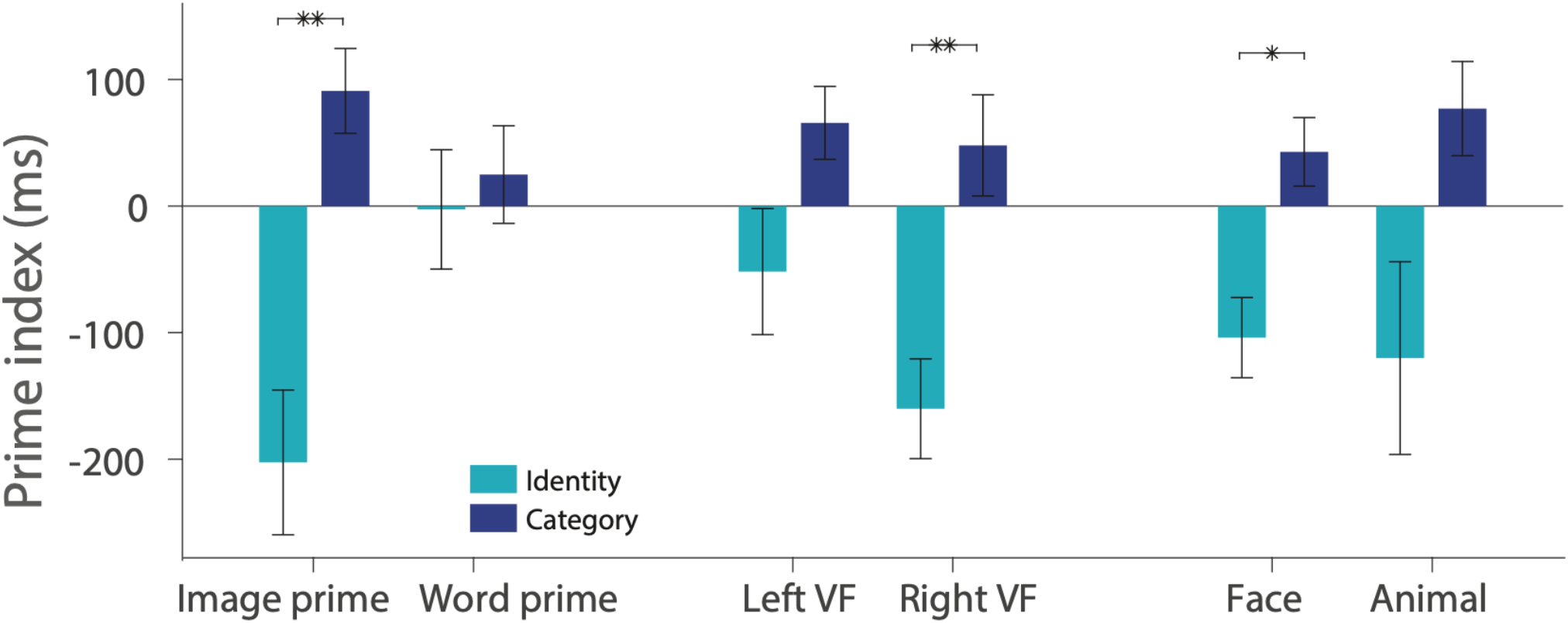
Influence of prime type, laterality, and target stimuli on priming effects. Additional analyses reveal priming effect variations across conditions. Effects are stronger for image primes than word primes, more pronounced in the right visual field (left-hemispheric processing), and greater for face targets than animal targets. Asterisks reflect Bonferroni–corrected p-values: *ns ≥ 0.05; * < 0.05; ** < 0.01*.

Binocular rivalry’s foveal presentation enabled us to test hemispheric differences. Conscious primes in right visual fields dissociated identity and category recognition that reveal left hemispheric bias (Figure 4; right visual field: Δ*PI*_*cat*−*id*_ = 205.0 ± 56.9 ms, *p*_*Bonf*_ = 0.009). In the left visual field, although the data exhibited the same trend observed elsewhere, the difference between identity and category recognition did not reach statistical significance. (Figure 4: left visual field: Δ*PI*_*cat*−*id*_ = 117.4 ± 57.7 ms, *p*_*Bonf*_ = 0.2).

Target stimuli also modulated priming. Face targets produced a significant dissociation (Figure 4; face: Δ*PI*_*cat*−*id*_ = 146.7 ± 45.0 ms, *p*_*Bonf*_ = 0.029), outperforming animal targets (Figure 4; animal: Δ*PI*_*cat*−*id*_ = 197.1 ± 88.5 ms, *p*_*Bonf*_ = 0.2;), though both showed the inversion pattern (faster identity, slower category RTs).

### Priming after effect depends on conscious awareness

Under unconscious priming, there was no sign of dissociation between identification and categorization (Figure 5A: Δ*PI*_*cat*−*id*_ = −14.2 ± 27.0 ms, *p* = 0.520), with effects absent across all conditions; image primes (Figure 5B: Δ*PI*_*cat*−*id*_ = 18.7 ± 41.3 ms, *p* = 0.380), word primes (Figure 5B: ΔPI_*cat*−*id*_ = −43.1 ± 56.2 ms, p = 0.420), left visual field (Δ*PI*_*cat*−*id*_ = 14.0 ± 54.6 ms, p = 0.800), right visual field (Δ*PI*_*cat*−*id*_ = −41.9 ± 57.2 ms, *p = 0.560)*, face targets (Δ*PI*_*cat*−*id*_ = 10.9 ± 37.5 ms, p = 0.800), and animal targets (Δ*PI*_*cat*−*id*_ = −38.4 ± 48.1 ms, p = 0.440). This absence aligns with reports linking certain recognition processes to conscious awareness (Amihai et al., 2011; Moradi et al., 2005), underscoring the condition-dependent nature of priming effects.

**Figure 5:**
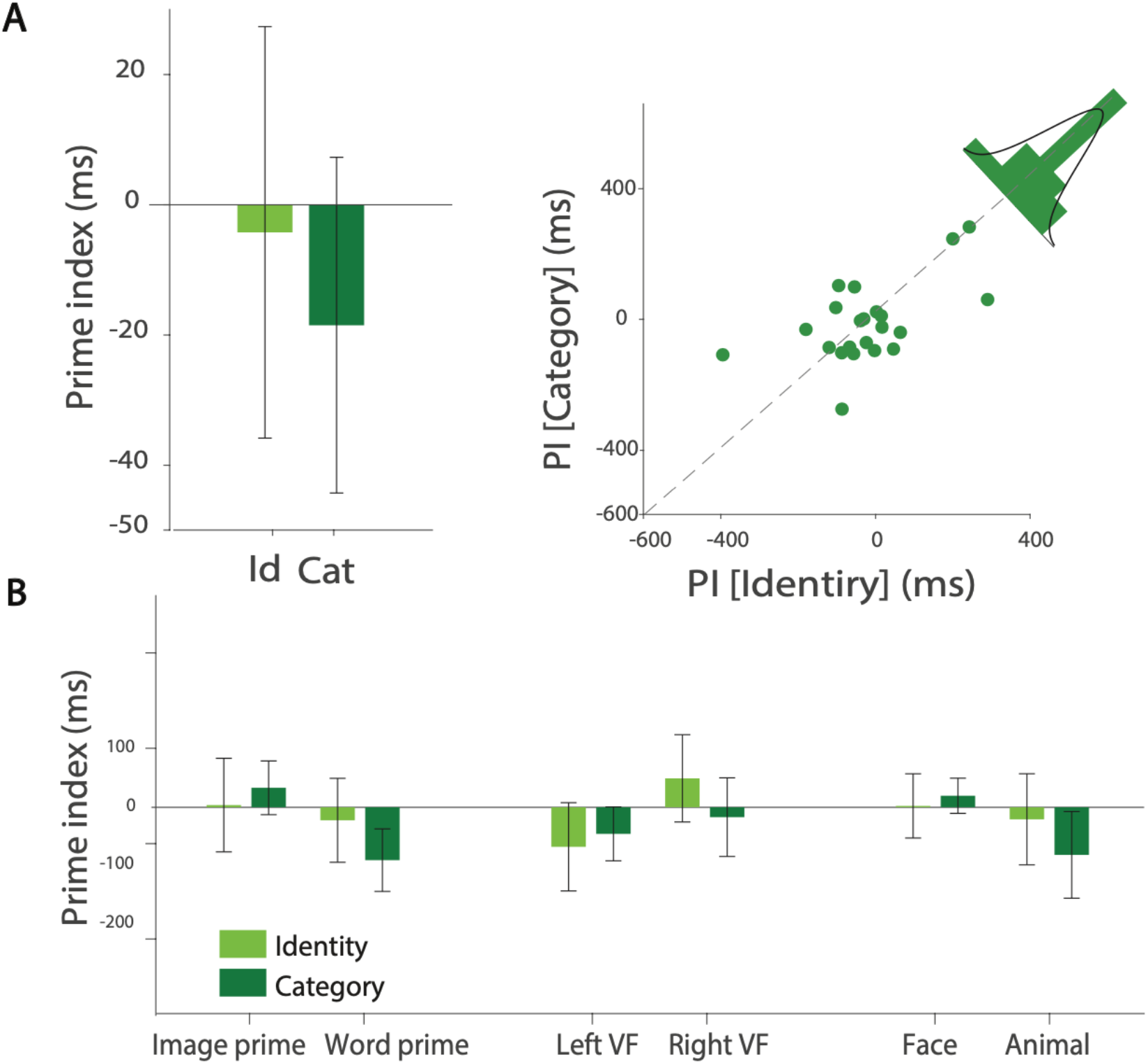
Unconscious priming effects on identity and category recognition. **(A)** Under unconscious priming, PI, scatter plot, and histogram show minimal variability between identity and category recognition, with no significant dissociation (specific PI values in text). **(B)** Priming effects across conditions (prime type, laterality, target stimuli) are not significant in unconscious trials. Error bars represent standard errors of the mean (SEMs) across participants.

### Modeling using DDM

We next present the model-based results from a drift-diffusion analysis of trial-wise decision and RTs. The model decomposes behavior into drift rate *υ* (efficiency of evidence accumulation), boundary separation a (response caution), starting point z (pre-decision bias), and non-decision time *T*_*er*_ (encoding/motor latency). We report how these components vary with priming (relevant vs. irrelevant), consciousness (conscious vs. unconscious), and recognition level (identity vs. category) to locate the source of the observed effects.

In all analyses we focused on within-participant difference scores contrasting trials with a relevant prime against those with an irrelevant prime. For each DDM parameter we computed Δ as the relevant minus irrelevant prime within each recognition level (identity/category) and consciousness condition: *Δv*(*rp* − *irp*) = *v*(*relevant prime*) − *v*(*irrelevant prime*); higher values indicate more efficient evidence accumulation under relevant primes, *Δa*(*rp* − *irp*) = *a*(*relevant prime*) − *a*(*irrelevant prime)*; positive values reflect greater response caution; *Δz*(*rp* − *irp*) = *z*(*relevant prime*) − *z*(*irrelevant prime)*; positive values indicate a pre-decisional bias toward the prime-consistent response; in our coding the upper boundary corresponds to ‘correct), and *ΔT*_*er*_(*rp* − *irp*) = *T*_*er*_ (*relevant prime*) − *T*_*er*_ (*irrelevant prime*); negative values indicate shorter non-decision latencies. Difference scores were estimated per participant and then tested at the group level.

### Conscious primes selectively enhance identity-level evidence

In the conscious priming condition, as shown in figure 6, drift rate was significantly higher for relevant compared with irrelevant primes during identity recognition (Figure 6A: Δ*v*(*rp* − *irp*)_*id*_ = 0.2 ± 0.04 ms, p < 0.001), indicating more efficient evidence accumulation for correct deciding. By contrast, the category task showed no reliable difference in drift rate between relevant and irrelevant prime (p = 0.3). This pattern implies that conscious primes selectively enhance identity-level evidence, predicting faster and more accurate identity decisions without a corresponding benefit for category decisions (Figure 6A: Δ*v*(*rp* − *irp*)_*cat*−*id*_ = 0.2 ± 0.05 ms, p < 0.001). during identity decision boundary separation (a) increased for relevant relative to irrelevant primes during identity judgments (Figure S3: Δ*a*(*rp* − *irp*)_*id*_ = 0.055 ± 0.02 ms; p = 0.03), consistent with a shift toward a more cautious response policy. No reliable change in a emerged for category judgments (Figure S3: Δa(rp − irp)_*cat*_ ≈ 0; p = 0.10), and the task-by-priming contrast (Identity vs. Category) was not significant (Figure S3: Δ*a*(*rp* − *irp*)_*cat*−*id*_ = 0.01 ± 0.03 ms; p = 0.60), indicating no evidence that the prime-induced change in response policy differs between tasks. In the same conscious priming condition, non-decision time showed no reliable change for relevant vs. irrelevant primes in either task (Figure S4: Δ*T*_*er*_(*rp* − *irp*)_*id*_ *p* = 0.8; Δ*T*_*er*_(*rp* − *irp*)_*cat*_ p = 0.8), and the task-by-priming contrast was also non-significant (Figure S4: Δ*T*_*er*_(*rp* − *irp*)_*cat*−*id*_ = 0.004 ± 0.01 ms; p = 0.8), indicating that priming did not measurably alter perceptual-encoding or motor latencies; the observed effects are better attributed to decision-stage dynamics (drift/response policy) rather than non-decision components.

**Figure 6:**
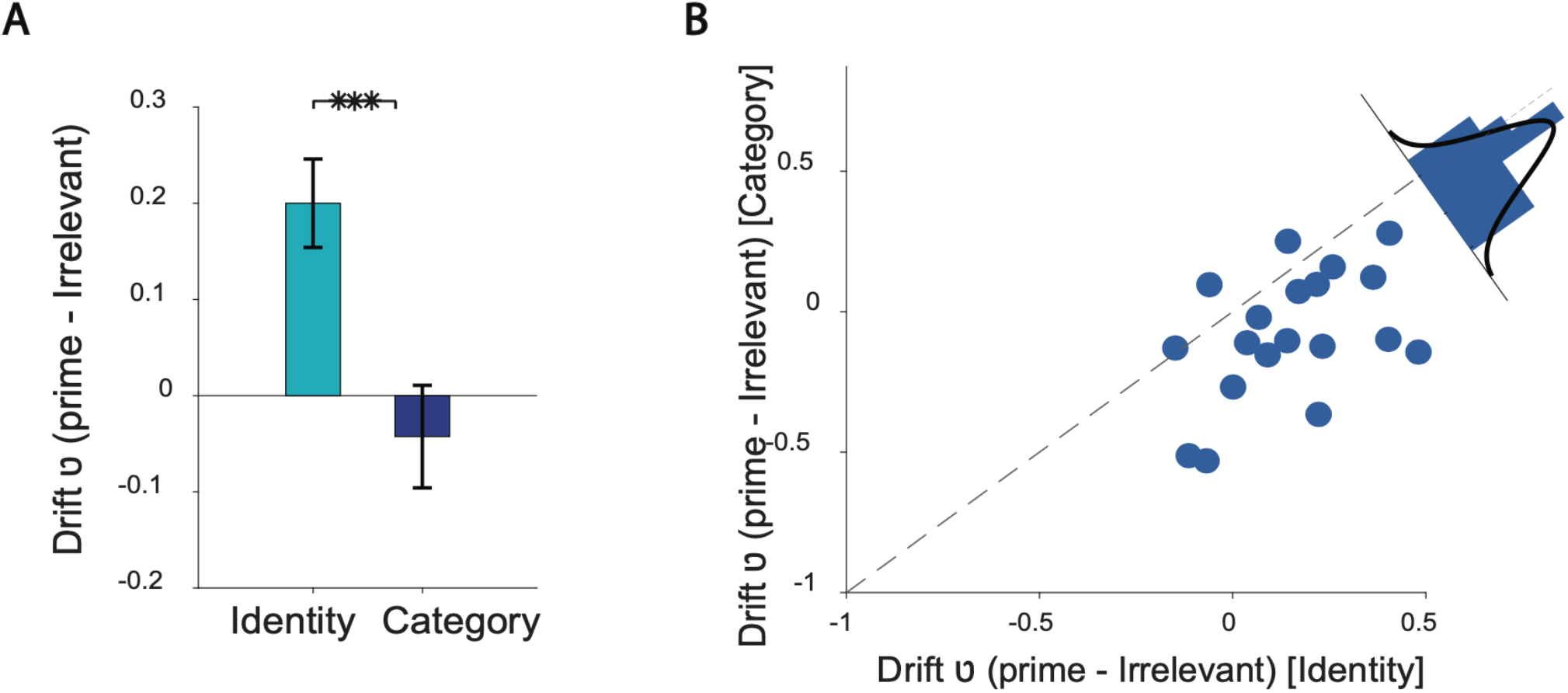
Drift rate (*v*) under conscious priming. **(A)** Group-level estimates of *υ* for relevant vs irrelevant primes are shown separately for identity and category recognition. Error bars represent standard errors of the mean (SEMs) across participants. Asterisks denote p-values for the relevant–irrelevant contrast within each recognition level: ***\*\*\**** *< 0.001*. **(B)** Scatter plot and histogram depict inter-subject variability. Points indicate participant-level estimates.

The starting point, in the conscious priming condition, exhibited a task-selective bias: for identity recognition, relevant primes shifted z away from an unbiased start (Figure 7A: Δ*z*(*rp* − *irp*)_*id*_ = 0.01 ± 0.008 ms; p = 0.04), indicating a pre-decisional tendency toward the prime-consistent response. No comparable shift was observed for category (Figure 7A: p = 0.1), and the identity– category contrast was significant (Figure 7A: Δ*z*(*rp* − *irp*)_*cat*−*id*_ = 0.03 ± 0.01 ms; p = 0.02). Together with the drift findings, these results suggest that conscious priming facilitates identity decisions through both enhanced evidence quality (higher *v*) and a starting-point bias, whereas category decisions show neither effect.

**Figure 7.**
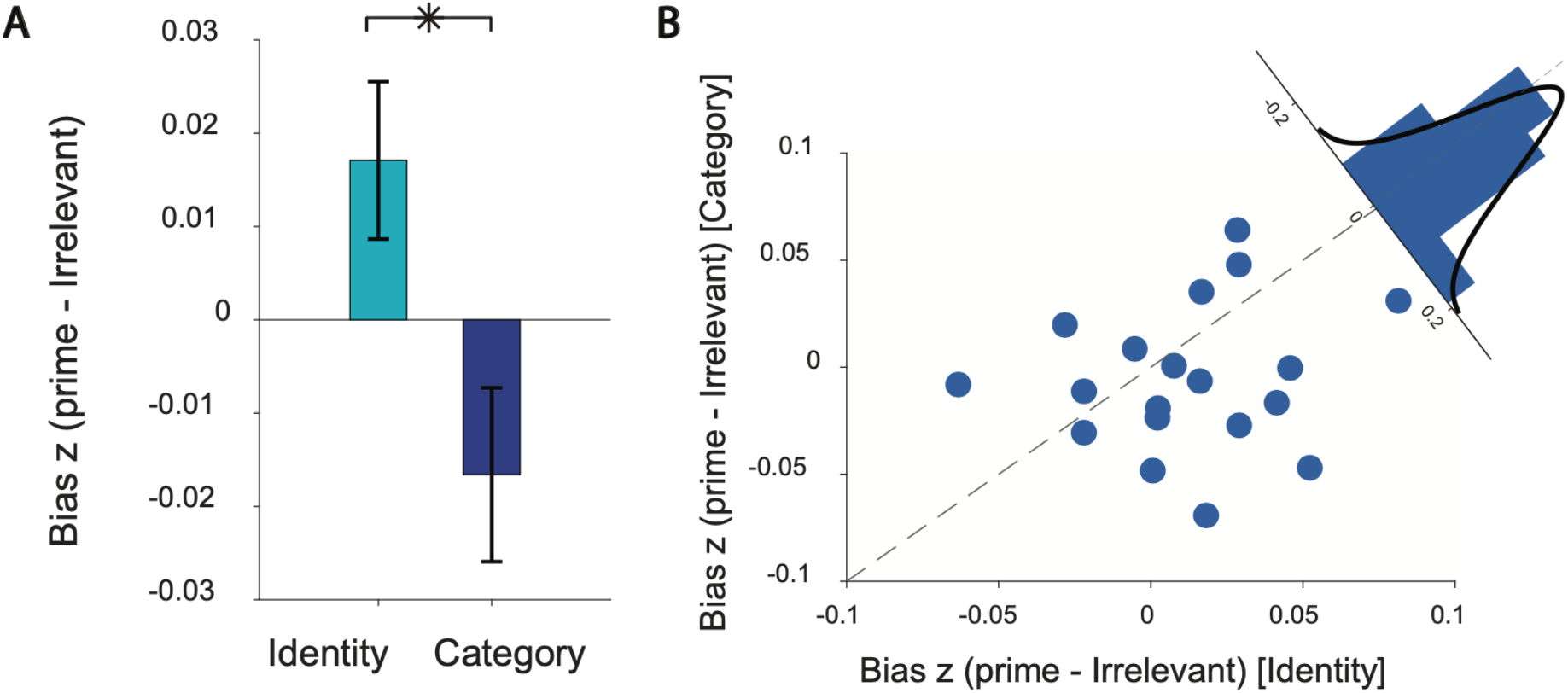
Starting point (z) under conscious priming. **(A)** Within-participant differences *Δz(rp-irp)* are plotted separately for identity and category decision. Positive values denote a bias toward the prime-consistent response (upper boundary = “Correct” under our coding). Identity showed a significant shift away from no-bias (p = 0.04), whereas category did not (p = 0.1). The between-task contrast was significant (Δz(rp − irp)_cat−id_ p = 0.02). **(B)** scatter plot and histogram depict inter-subject variability. Points indicate participant-level estimates.

Drift–diffusion modeling showed that under conscious priming, identity decision—but not category—exhibited higher drift and a starting-point bias toward the relevant prime response, yielding faster and more accurate identity decisions. Boundary separation was modestly elevated for identity (with no reliable change for category), while non-decision time was unchanged, indicating that the effects arise primarily from decision-stage dynamics rather than encoding or motor latencies.

In the unconscious priming condition, none of the DDM parameters showed significant differences between related and unrelated primes across identity and category judgments (Δ*v*(*rp* − *irp*)_*cat*−*id*_ = 0.07 ± 0.08 ms; p = 0.6, Δ*a*(*rp* − *irp*)_*cat*−*id*_ = 0.03 ± 0.05 ms; p = 0.6, Δ*T*_*er*_(*rp* − *irp*)_*cat*−*id*_ = −0.006 ± 0.01 ms; p = 0.4, Δ*z*(*rp* − *irp*)_*cat*−*id*_ = −0.006 ± 0.01 ms; p = 0.8), confirming that unconscious primes did not measurably modulate evidence accumulation, decision boundaries, non-decision times, or starting biases.

## Discussion

The present study reveals a striking dissociation in how visual priming affects recognition at different levels of abstraction (identification and categorization). Across all conditions, the basic-level (category) advantage remained robust, with category judgments being faster than identity judgments overall, consistent with prior work on hierarchical object recognition (Rosch et al., 1976). A particularly novel aspect of our findings is the opposite direction of priming effects under conscious priming. Our data reveal that consciously perceived primes not only facilitate identity recognition but also slowed down category recognition. This suggests that conscious priming amplifies detailed analysis of specific features (e.g., individual men) but disrupts rapid grouping into broader categories (e.g., men vs. women).

Image primes proved more effective than word primes, particularly for identity judgments, underscoring the advantage of visual over verbal information in supporting fine-grained recognition. This pattern is consistent with prior evidence that visual cues facilitate subordinate-level identification more efficiently than linguistic labels (Carr et al., 1982; Reinitz et al., 1989). Furthermore, within the visual modality, the benefit of image primes was especially pronounced for faces, aligning with extensive literature on the privileged role of facial features in perception and recognition(Gao et al., 2022; Zhou et al., 2010). Together, these findings emphasize both a general efficiency of visual information for identity recognition and the specialized neural mechanisms dedicated to face processing. A left-hemispheric bias emerged in conscious priming, likely reflecting specialized cortical areas for detailed visual and linguistic processing (Koivisto & Revonsuo, 2000; Marsolek, 1999). In contrast, identity-category divergence did not emerge under unconscious priming, highlighting that the presence of conscious awareness changes the way primes interact with recognition processes. Interpreted in the context of drift diffusion decision-making models (Ratcliff & McKoon, 2008; Voss et al., 2013), these results suggest that conscious primes selectively enhance evidence accumulation when the task demands fine-grained identification, but introduce impairment when the task requires broader categorical judgments.

Consciously perceived primes facilitated identity recognition while impairing category recognition, whereas unconscious primes produced only a nonsignificant facilitation of category judgments. This pattern is consistent with a fine-to-coarse account of visual processing, in which conscious awareness sharpens fine-grained, subordinate-level representations, enhancing access to individual identities but disrupting rapid basic-level categorization that depends on coarser information(Dehaqani et al., 2016; Koivisto & Rientamo, 2016). These findings emphasize that conscious and unconscious priming differ not only in magnitude but within the neural machinery, both feedforward and feedback processes contribute to the representation of information across different levels of abstraction (Chien et al., 2023; Stein et al., 2020).

Traditional analyses of priming rely on comparing mean reaction times or accuracy across conditions. While such analyses revealed the high-level pattern (speeding vs. slowing), they leave ambiguous *how* the prime exerts its influence. By applying the DDM to our data, we were able to attribute the effects of priming to specific latent parameters associated with decision-making. This affords a more precise interpretation. For example, the finding that conscious identity primes elevated the drift rate but did not reduce the non-decision time indicates that the primes likely improved the quality of perceptual evidence or decision-level information, rather than simply speeding up early perceptual processing or motor execution. In other words, participants were not just pressing the button faster; they were genuinely accumulating information more rapidly towards the correct decision when a relevant prime was visible.

Our data reinforce that the unconscious primes did not modify drift rates, caution, starting biases, or non-decision times – in other words, they did not measurably enter into the decision computations. Consistent with prior behavioral studies, truly subliminal primes (especially under CFS) might not provide a strong enough or sustained enough signal to influence decision outcomes on a single-trial basis. Stein and colleagues (Stein et al., 2020), for instance, reported that pictures suppressed by CFS produced at best weak semantic priming effects, much smaller than those obtained with backward masking paradigms. Chien and colleagues (Chien et al., 2023) concluded that unconscious semantic activation might not propagate far enough to affect decision-making unless the conditions are highly optimized. Our results are in alignment: under our controlled but stringent conditions, unconscious primes were effectively inert.

Our findings also support the robust basic-level advantage, a hallmark of the categorization literature (Jolicoeur et al., 1984; Rosch et al., 1976). Prior research has suggested that with increasing familiarity or expertise, processing at the subordinate level can functionally shift to the basic level, such that exemplars once recognized slowly become the most efficient entry point for categorization(Johnson & Mervis, 1997; Tanaka & Taylor, 1991). In our paradigm, the presence of a conscious prime appears to mimic this mechanism by supplying additional information that enhances subordinate recognition, thereby attenuating the basic-level advantage. What is particularly intriguing, however, is that no prior work has suggested a comparable transformation at the basic level itself. Our results raise the possibility that when primes provide excessive information, basic-level judgments may in effect become similar to the superordinate level. In such cases, categorization requires additional integration time, producing relative slowing despite the ordinarily privileged status of basic-level categories. While prior studies have shown that subordinate recognition improves with additional information or training (e.g., through high spatial frequency details or expertise), they did not examine a simultaneous cost to basic-level categorization. Our findings suggest that priming can create precisely such a cost, revealing a novel dynamic in the hierarchy of visual categorization.

Such an opposite-direction effect within a single experiment is, to our knowledge, unprecedented in prior work and therefore represents a novel empirical contribution. It suggests that priming is not a uniformly beneficial process – the outcome depends on an interaction between what information the prime provides and what the decision requires. Drift Diffusion Modeling (DDM) provided insight into the underlying mechanisms: under conscious priming, the drift rate (evidence accumulation speed) increased for identity decisions but not for category decisions. An elevated drift rate indicates more efficient accumulation of decision evidence, explaining the speed-up in identity recognition. Notably, conscious identity priming also induced a small but significant starting-point bias in favor of the prime-consistent response. This suggests that when the prime was visible, participants entered the decision with a pre-existing push toward confirming the primed identity, leading to faster (and potentially more accurate) identity judgments.

Under unconscious priming conditions, DDM parameters (drift, bias, caution, decision threshold) remained essentially unchanged, indicating that unseen primes failed to modulate the decision process. Together, these findings highlight a double dissociation: conscious awareness is required for priming to exert robust effects, and those effects can be facilitatory or impairing depending on the level of recognition. This dissociation is a key contribution of our work, underscoring the importance of considering both the level of abstraction (identity vs. category) and conscious access in understanding priming phenomena.

We encourage future research on conscious and unconscious priming to adopt similar approaches. By combining sophisticated behavioral modeling with neuroscience methods (e.g., EEG/MEG or neural network modeling), one could test whether the drift rate increases we observed correspond to neural signatures of enhanced sensory evidence or higher signal-to-noise ratio in cortical areas, and whether the starting bias correlates with pre-stimulus brain states or attentional orienting triggered by the prime.

