## supplemental figures for "Opposite Priming Effects on Identity vs. Category Recognition Require Conscious Awareness"

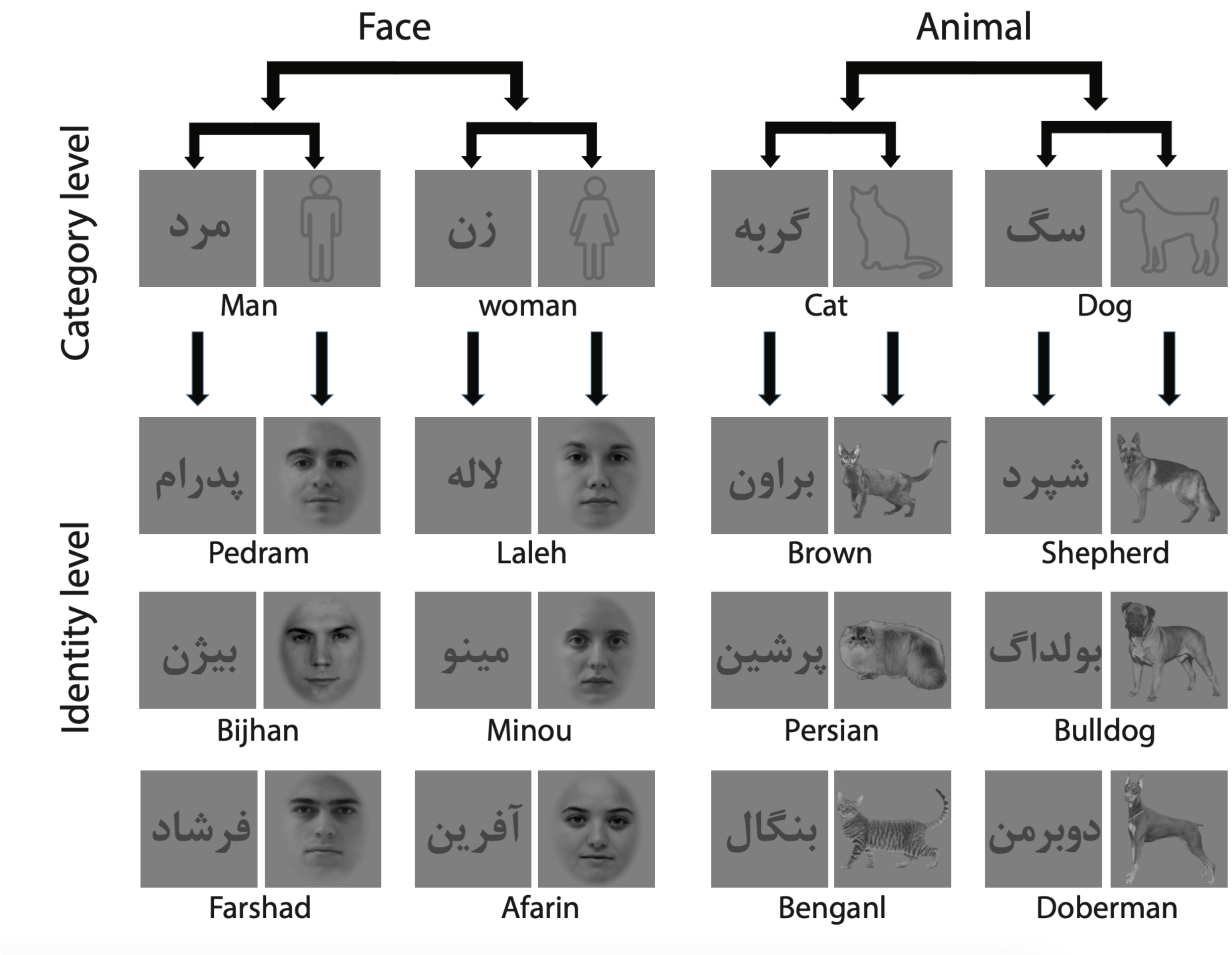


**Figure S1. Stimulus set**

The hierarchy of stimuli and the stimuli presented in the experiment are outlined as follows. Stimuli were presented at two levels of abstraction: categorization (man/woman and cat/dog) and identification within each of the four groups, with three stimuli for each identification task. The specific stimuli in the four groups were:

Woman: مینو (Minou), آفرین (Afarin), لاله (Laleh)
Man: فرشاد (Farshad),بیژن (Bijhan), پدرام (Pedram)
Cat: بنگال (Bengal), پرشین (Persian), براون (Brown)
Dog: شپرد(Shepherd), دوبرمن (Doberman), بولداگ (Bulldog)

Stimuli were presented both as images and written words. Given that the subjects were Iranian, the language used for the written words was Persian.

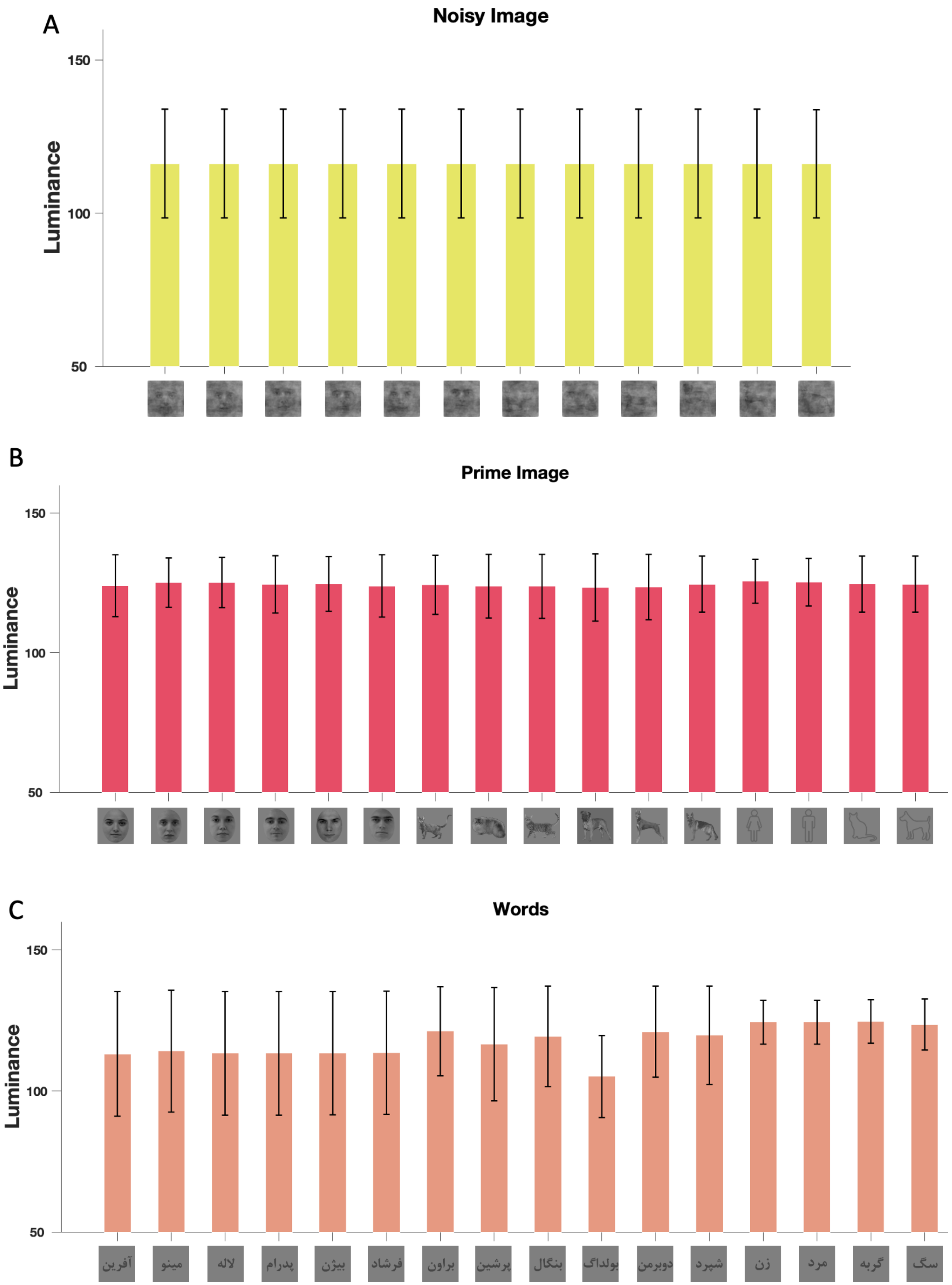
A
Noisy Image
Luminance
150
100
50
B
Prime Image
Luminance
150
100
50
C
Words
Luminance
150
100
50
آفرین
مینو
لاله
پدرام
بیژن
فرشاد
براون
پرشین
بنگال
بولداگ
دوبرمن
شپرد
زن
مرد
گربه
سگ

**Figure S2. Similarity of stimuli's luminance**

The figure illustrates the luminance levels of all stimuli used in the experiment. Panel (A) displays noisy images utilized for target stimuli, Panel (B) shows prime images used for relevant and irrelevant prime stimuli, and Panel (C) presents words used for category names and word primes. Error bars in the figures represent the Standard Errors of the Means (SEMs) calculated across subjects.

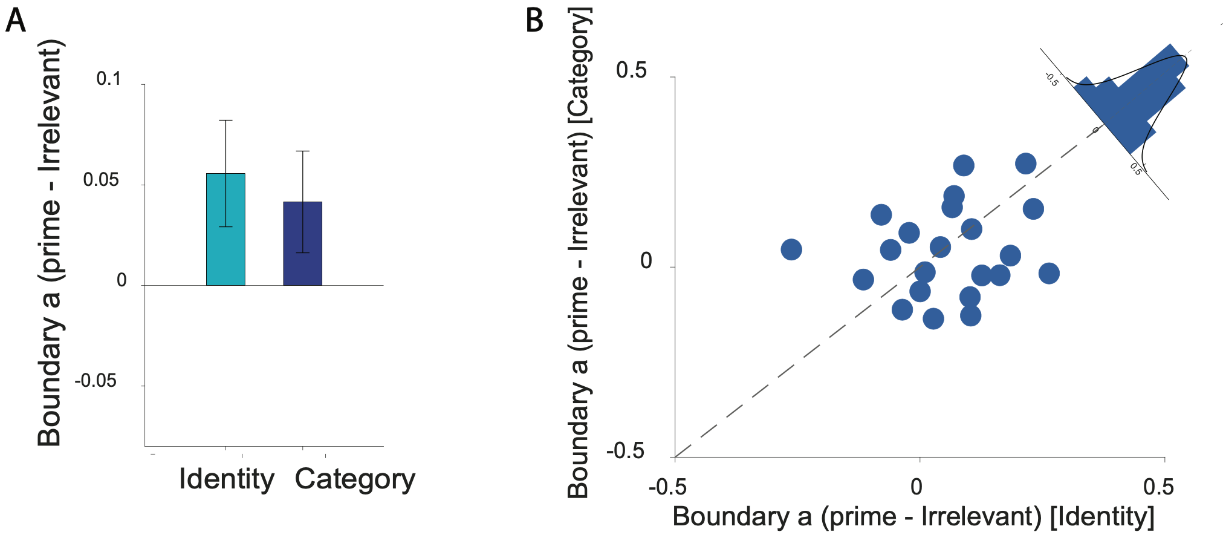


**Figure S3. Boundary separation (a) under conscious priming.**

(A) Within-participant differences Δa(pr-ir) are shown separately for Identity and Category judgments. A significant positive shift was observed for Identity ($p = 0.03$), whereas Category showed no reliable effect ($p = 0.10$). The difference between tasks (Identity vs. Category) was not significant ($p = 0.6$). (B) scatter plot and histogram depict inter-subject variability. Points indicate participant-level estimates.

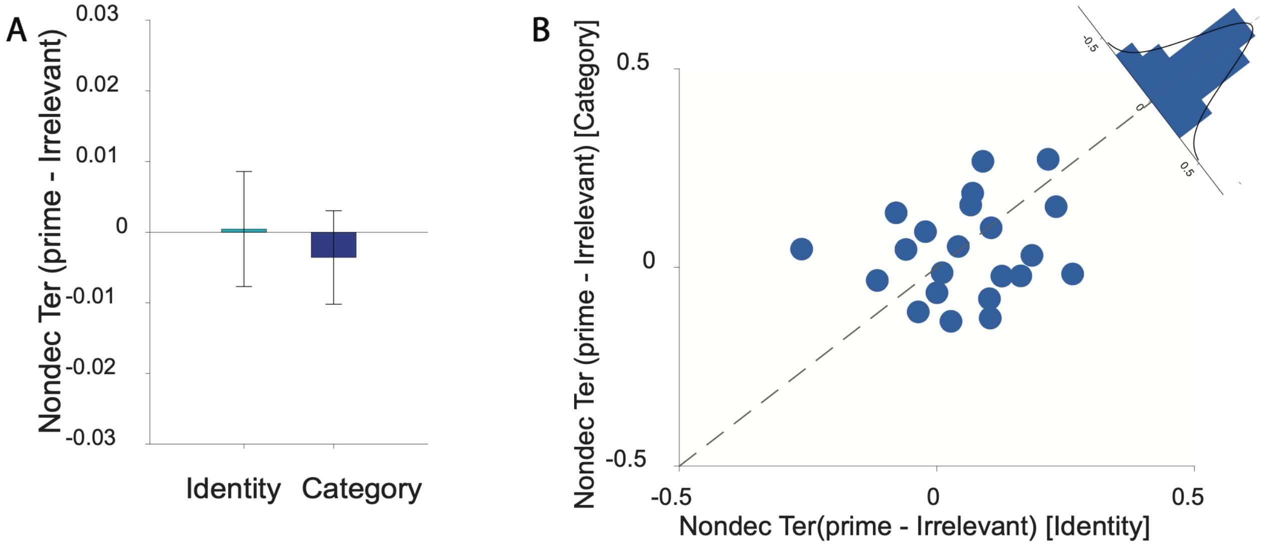


**Figure S4. Non-decision time ($T_{er}$) under conscious priming.**

Within-participant differences $\Delta T_e$ (pr-ir) are shown separately for Identity and Category judgments Neither task differed reliably from zero (Identity: ns; Category: ns), and the Identity–Category contrast was also non-significant (ns). (B) scatter plot and histogram depict inter-subject variability. Points indicate participant-level estimates.
